# High glucose levels impair the body’s renal protective function during water deprivation by blunting the FXR-TonEBP axis

**DOI:** 10.1101/2025.07.11.664291

**Authors:** Tuo Wei, Jiebo Huang, Yan Li, Liru Yin, Fang Tian, Qiong Wang, Enchao Zhou

## Abstract

**Background:** Dehydration often leads to kidney-related complications, yet its impact on diabetic patients remains underexplored. This study aimed to investigate the effects of dehydration on diabetic nephropathy(DN) and the underlying mechanisms of injury.

**Methods:** We analyzed clinical outcomes in 81 dehydration-induced acute kidney injury patients (35 with diabetes) and validated findings in streptozotocin-induced diabetic mice subjected to intermittent water deprivation (WD). Renal function, histopathology, and molecular markers (FXR, TonEBP, AQP2, gp91, apoptosis regulators) were assessed.

**Results:** Kidney injury was aggravated after water deprivation in patients with diabetics and it is difficult to recover after water supplementation, with poor prognosis. Animal experiments have shown that compared with mice in a normal state, dehydration challenge aggravated renal damage in diabetic mice with increased serum creatinine, pathological damage, and renal interstitial cell apoptosis. Mechanistically, in a normal state FXR-TonEBP axis was enhanced to protect the kidney after water deprivation challenge, but this compensatory protective response was blunted in the diabetic state.

**Conclusion:** Hyperglycemia blunts the FXR-TonEBP-osmolyte pathway during dehydration, compromising renal medullary protection. Targeting this axis may mitigate dehydration risks in diabetic nephropathy.

## 1. Introduction

Dehydration is a common clinical problem that frequently causes a hypertonic state in the body, leading to damage in multiple organs, particularly the kidneys ^[1, 2]^. Numerous factors may induce dehydration, including insufficient fluid consumption, profuse sweating, vomiting, diarrhea, and excessive diuretic use ^[3]^. Prior research has recognized certain high-risk groups for dehydration, including young children, older adults, athletes, and individuals working outdoors in high temperatures ^[4-6]^. Nevertheless, the potential for dehydration-related renal injury in patients with pre-existing diseases, especially those with diabetes, has not been sufficiently investigated.

Dehydration impairs renal function through multiple pathways. First, dehydration-induced hypertonicity triggers elevated vasopressin (copeptin) secretion, leading to renal tubular injury and oxidative stress in the kidneys^[6]^. Furthermore, dehydration stimulates the polyol metabolic pathway, where hyperosmolality increases aldose reductase activity, promoting glucose conversion into sorbitol. As an osmotic agent, sorbitol helps protect renal tubular cells and medullary interstitial cells from hypertonic damage while enhancing water reabsorption.

The vasopressin and aldose reductase pathways activated during dehydration are typically considered protective mechanisms, as they promote urine concentration during water scarcity^[7]^. However, emerging evidence suggests that chronic activation of these pathways may contribute to progressive renal injury^[8]^. We propose that such damage could be amplified in hyperglycemic conditions, where increased glucose concentrations may further elevate renal medullary hyperosmolarity^[9, 10]^.

Dehydration impairs renal function through multiple pathways. The primary mechanism involves dehydration-induced hypertonicity, which elevates vasopressin (copeptin) levels, subsequently causing renal tubular injury and oxidative stress in renal tissues^[11, 12]^. Renal medullary hypertonicity drives high TonEBP expression in medullary epithelial cells. Elevated TonEBP enhances urine concentration by upregulating aquaporin 2 (AQP2) expression, thereby protecting the renal medulla from osmotic damage.The farnesoid X receptor (FXR), a ligand-activated transcription factor, contributes to water balance regulation by increasing AQP2 expression in renal medullary collecting ducts^[13]^. Emerging evidence demonstrates that FXR promotes both expression and nuclear translocation of TonEBP, along with its downstream targets including the sodium-myo-inositol transporter (SMIT) and heat shock protein 70 (HSP70). This FXR-TonEBP signaling cascade provides cellular protection for renal medullary interstitial cells during hyperosmotic stress^[14]^.

In summary, dehydration exerts detrimental effects on renal medullary function, a process mediated by the FXR-TonEBP signaling pathway ^[15-17]^. Nevertheless, whether this regulatory mechanism differs in patients with underlying pathologies, particularly DM, remains unclear. Could the expression patterns of FXR and TonEBP be modified under such pathological conditions? Addressing these critical questions constitutes the primary objective of our investigation.

## 2. Materials and methods

### 2.1. Patients’ medical and clinical data

This retrospective study analyzed the medical records of 81 patients (49 males and 32 females) diagnosed with abnormal renal function. Data were collected from standard clinical examinations conducted between December 2019 and November 2024, with no additional risks imposed on participants. The Ethics Committee of Nanjing Pukou District Hospital of Traditional Chinese Medicine granted a waiver for informed consent (Approval No. 20210027) as the study utilized anonymized retrospective data.

Inclusion criteria comprised: (1) no prior history of renal disease, and (2) presence of acute kidney dysfunction (eGFR <90 mL/min/1.73 m^2^) attributable to dehydration (including diarrhea, excessive sweating, or inadequate fluid intake). Demographic information, hospitalization timelines, underlying comorbidities, and laboratory parameters (serum urea nitrogen and creatinine) were recorded at admission and post-treatment.

Patients were followed for four months, with renal recovery outcomes classified as: complete recovery (normalization of creatinine levels), partial recovery (reduced but persistently elevated creatinine), or no recovery (increased creatinine versus baseline).

### 2.2. Animal models and treatment protocols

Ethical Approval and Experimental Animals:All animal experiments were approved by the Experimental Animal Ethics Committee of Nanjing Pukou District Hospital of Traditional Chinese Medicine (Approval No. 20210026) and strictly followed NIH guidelines for laboratory animal care. twenty male C57BL/6 mice (8-9 weeks old, 19-21 g) from Zhejiang Weitonglihua Laboratory Animal Technology Co., Ltd. were housed under controlled conditions (12-h light/dark cycle, 22±1°C) with free access to food and water. After one-week acclimatization, mice were randomly divided into four groups (n=5/group): blank control (BC), BC with water deprivation (BC+WD), diabetes mellitus (DM), and DM with water deprivation (DM+WD).

Diabetes Induction and Experimental Protocol:DM groups received daily intraperitoneal STZ injections (50 mg/kg in 100 mM sodium citrate buffer, pH 4.5) for 5 consecutive days, while controls received citrate buffer alone. Diabetes was confirmed by fasting blood glucose >16.7 mmol/L one week post-injection. All mice were fed a Western diet, with WD groups subjected to 24-h water deprivation every other day for 16 weeks. Physiological parameters (body weight, water intake, urine output, blood glucose) were monitored biweekly.

Sample Collection and Processing:After 16 weeks, blood samples were collected via cardiac puncture and centrifuged (4°C, 3000 rpm, 10 min) for serum separation. Kidneys were processed for: (1) left kidneys fixed in 4% paraformaldehyde for histopathology and immunofluorescence, and (2) right kidneys snap-frozen in liquid nitrogen for molecular analyses.

### 2.3. Blood and urine chemistry

To evaluate blood and urine biochemistry parameters, the animals were housed in metabolic cages for 24 hours to facilitate urine sample collection. Urinary protein levels were quantified over the 24-hour period using a protein analyzer (BIOSTEC, Spain, model BA400). Renal function was evaluated by analyzing blood serum samples with an automated analyzer (Hitachi, Tokyo, Japan, model 7180).

### 2.4. Reagents

The following primary antibodies were employed: anti-FXR (Proteintech, Wuhan, China, no.25055-1-AP), anti-TonEBP (Affinity Biosciences, Cincinnati, OH, USA, no.AF7663), anti-caspase-3 (Cell Signaling Technology, Shanghai, China, no.9662), anti-Bcl-2 (Proteintech, Wuhan, China, no.26593-1-AP), anti-BAX (Proteintech, Wuhan, China, no.50599-2Ig), anti-gp91 (BD Biosciences, Santa Clara, CA, USA, no.611414), anti-AQP2 (Abclonal, Wuhan, China, no.A16209) and anti-β –actin (Proteintech, no.66009-1-Ig). Secondary antibodies consisted of anti-mouse IgG (H+L) (Proteintech, no.SA00001-1) and anti-rabbit IgG (H+L) (Proteintech, no.SA00001-2).

### 2.5. Western blot analysis

Kidney homogenates were prepared by lysing renal cells in a buffer containing protease and phosphatase inhibitors (Beyotime, China). The lysates were sonicated and centrifuged to obtain the supernatant, which was then mixed with SDS and heated at 100°C for 10 min for protein denaturation. Protein separation was performed using 10% SDS-PAGE (Bio-Rad, China), followed by transfer to PVDF membranes (Millipore, USA). The membranes were blocked with 5% non-fat milk in PBST (PBS with 0.1% Tween-20) for 1h at room temperature. Primary antibody incubation was carried out overnight at 4 ° C, followed by secondary antibody incubation for 1 h at room temperature. After washing, protein bands were detected using an HRP chemiluminescent substrate and visualized with a ChemiDoc™ XRS system (Bio-Rad, USA). Band intensity was quantified using ImageJ software (v1.52a).

### 2.6. Histopathology

Kidney tissues were fixed in 4% paraformaldehyde, dehydrated in graded alcohols, and subsequently stained with hematoxylin-eosin (Servicebio, Wuhan, China, no.GP1031), Masson’s trichrome (Servicebio, no.GP1032), and Periodic Acid-Schiff (PAS, Servicebio, no.GP1039). Histological analysis was performed using ImageJ software for quantitative assessment of stained areas.

### 2.7. Immunofluorescence staining

Renal tissues were sectioned at 40 µm thickness using a microtome (Leica RM2016, Shanghai, China). Free-floating sections were processed for immunofluorescence staining with primary antibodies against FXR, AQP2, TonEBP, and gp91. Stained sections were examined under a microscope (Nikon eclipse ci-e, Japan) and imaged at 10× magnification using a digital slide scanner (3DHISTECH Pannoramic MIDI, Hungary). Quantitative analysis of immunoreactivity was performed using ImageJ software, with antibody expression levels expressed as integrated density values.

### 2.8. Tunel assay

Apoptotic renal tissue cells were labeled using a terminal deoxynucleotidyl transferase dUTP nick end labeling (Tunel) kit (Roche Diagnostics, Indianapolis, USA, Cat. no.11684817910) according to the manufacturer’s protocol. Quantification was performed under an immunofluorescence microscope (Nikon, Japan, ECLIPSE Ci Series). Five randomly selected fields (×200 magnification) were analyzed to determine the proportion of apoptotic cells relative to the total cell count. ImageJ software was employed to measure the stained areas of apoptotic cells.

### 2.9. Statistical data evaluation

Statistical analyses were performed using GraphPad Prism 7, with data presented as mean ± SD. Normality of distribution was assessed using the Shapiro-Wilk test. For comparisons between two groups, an unpaired t-test was employed, while one-way ANOVA was used for multi-group comparisons. A p-value <0.05 was considered statistically significant.

## 3. RESULTS

### 3.1 Among the patients with renal injury caused by water deprivation, those with DM have the worst prognosis

Demographic and medical data of patients with renal injury caused by dehydration are displayed in Table 1. In this study, older people were defined as those over 65 years of age. Our analysis of these data showed that 47 individuals (58%) were older than 65 years whereas 34 (42%)were younger. 49 patients (60.5%) were males and 32 (39.5%) were females. The reported main causes of dehydration among these 81 patients were Poor appetite in 31 individuals (38.3%), diarrhoea in 24 (29.6%), vomit in 10 (12.4%), hyperpyrexia in 9 (11.1%), drinking water disorders in 5 (6.2%) and excessive exercise in 2 (2.5%). A total of 67 patients(80.3%) had chronic diseases, including 44(54.3%) with hypertension, 35(43.2%) with diabetes, and 16(19.8%) with cerebral infarction. Only 14(17.3%) patients had no chronic diseases.

**Table 1.** Demographic and medical data of patients with renal injury caused by dehydration.

| Variates | n=81 | Ratio(n%) |
| --- | --- | --- |
| Age |  |  |
| ≥65(years) | 47 | 58% |
| < 65(years) | 34 | 42% |
| Gender |  |  |
| Male | 49 | 60.5% |
| Female | 32 | 39.5% |
| Dehydration causes |  |  |
| Drinking water disorders | 5 | 6.2% |
| Poor appetite | 31 | 38.3% |
| Diarrhoea | 24 | 29.6% |
| Excessive exercise | 2 | 2.5% |
| Vomit | 10 | 12.4% |
| Hyperpyrexia | 9 | 11.1% |
| Underlying diseases* |  |  |
| Hypertension | 44 | 54.3% |
| Diabetes | 35 | 43.2% |
| Cerebral infarction | 16 | 19.8% |
| No underlying diseases | 14 | 17.3% |

\*Underlying diseases: The top three diseases were hypertension, DM and cerebral infarction.

eGFR at onset and prognosis in patients with different underlying diseases are displayed in Table 2. According to the degree of renal damage at the time of onset, eGFR less than 15ml/min was defined as severe damage, followed by DM (14 cases), hypertension (12 cases), cerebral infarction (3 cases), and no underlying disease (2 cases). However, we found that prognosis significantly varied among patients who had different comorbid chronic diseases. Data showed that all patients without chronic diseases recovered fully. By comparison, some of the patients with hypertension, DM not recover after treatment. respectively. Among the 7 patients with worsened prognosis, 6 (85.7%) had DM.These data proved that in a dehydrated state, diabetics displayed a relatively high the worst prognosis.

**Table 2.**
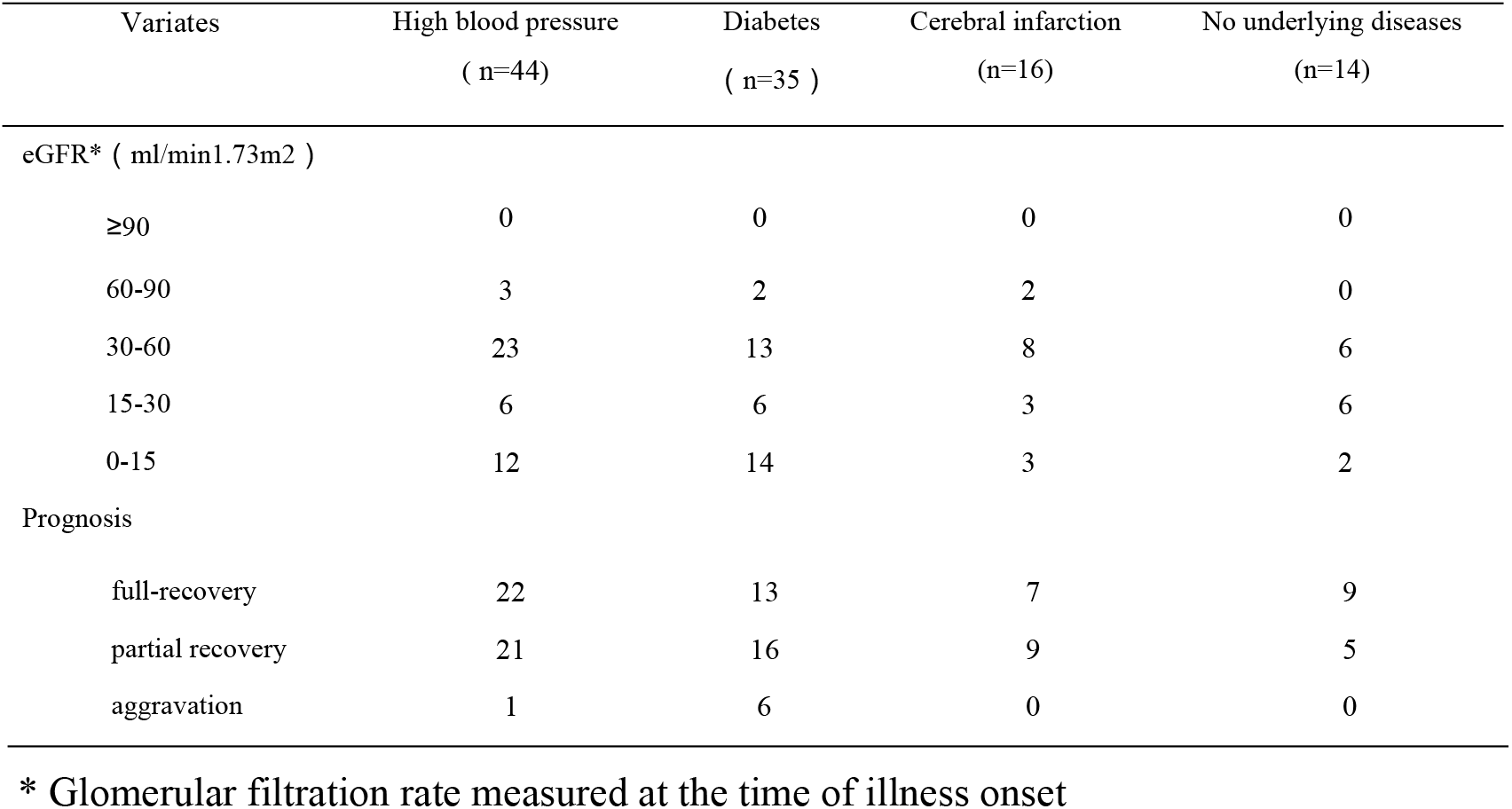
eGFR at onset and prognosis in patients with different underlying diseases.

| Variates | High blood pressure<br>( n=44) | Diabetes<br>( n=35 ) | Cerebral infarction<br>(n=16) | No underlying diseases<br>(n=14) |
| --- | --- | --- | --- | --- |
| eGFR* ( ml/min1.73m2 ) |  |  |  |  |
| ≥90 | 0 | 0 | 0 | 0 |
| 60-90 | 3 | 2 | 2 | 0 |
| 30-60 | 23 | 13 | 8 | 6 |
| 15-30 | 6 | 6 | 3 | 6 |
| 0-15 | 12 | 14 | 3 | 2 |
| Prognosis |  |  |  |  |
| full-recovery | 22 | 13 | 7 | 9 |
| partial recovery | 21 | 16 | 9 | 5 |
| aggravation | 1 | 6 | 0 | 0 |

\* Glomerular filtration rate measured at the time of illness onset

These results show that it is of utmost importance to further study the pathological changes and mechanisms underlying renal damage in response to dehydration. For this purpose, we established animal models for further research.

### 3.2 Kidney injury was exacerbated in diabetic mice following water deprivation

Compared to BC mice, DM mice exhibited characteristic diabetic features, including lower body weight and increased water intake (Figure 1A,B). Notably, under normal drinking conditions, DM+WD mice produced more urine and had higher urinary protein levels than DM mice, while no such differences were observed between BC and BC+WD mice (Figure 1C,D). Water deprivation did not significantly alter urea nitrogen or uric acid levels (Figure 1E,G). However, serum creatinine levels were higher in DM+WD mice than in DM mice, whereas BC mice remained unaffected (Figure 1F). Blood glucose levels in DM mice remained stable throughout the experiment (Figure 1H). Histopathological analysis further revealed renal damage in DM mice after water deprivation: while HE staining showed no significant differences between non-DM and DM groups (Figure 1I), PAS staining demonstrated tubular blockages and dilations in the renal collecting ducts of DM+WD mice (Figure 1J). Additionally, Masson staining showed that, compared with the BC, BC + WD, and DM groups, the DM + WD group exhibited more significant fibrosis(Figure 1K). These findings suggest that water deprivation exacerbates renal medullary damage in DM mice, impairing urine concentration and leading to polyuria, while renal fibrosis likely contributes to reduced filtration function, as evidenced by elevated serum creatinine levels.

**Figure 1.**
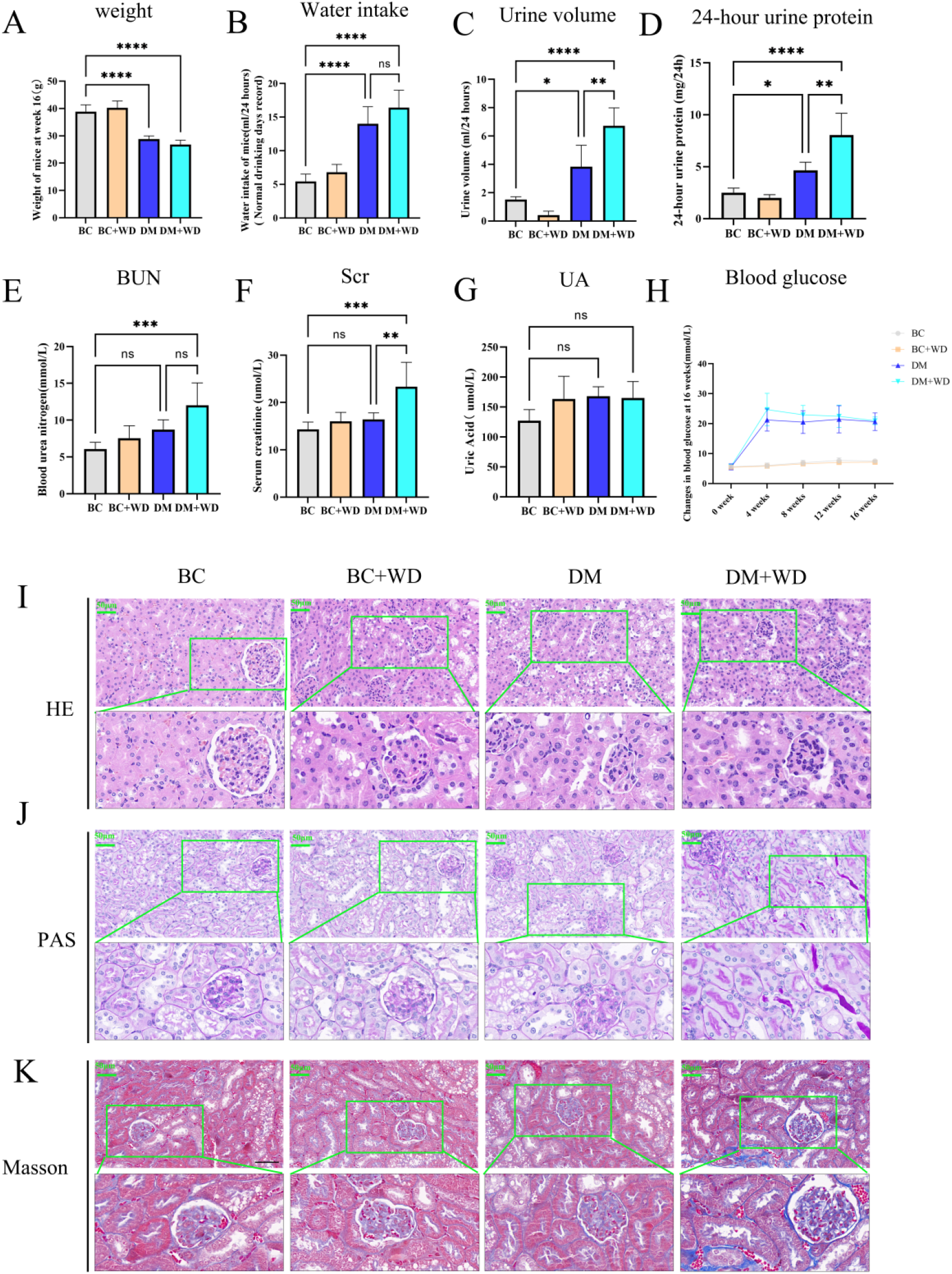
Metabolic alterations and renal histopathological changes in diabetic mice following water deprivation. (A) body weight. (B) water in take. (C) urine output(24h). (D) 24h urine protein. (E) blood urea nitrogen. (F) serum creatinine. (G)blood uric acid. (H) blood glucose levels(16 weeks). (I) HE staining of renal tissue(Scale 50 μm). (J) PAS staining of renal tissue(Scale 50 μm). (K) Masson staining of renal tissue(Scale 50 μm). Data are presented as mean ± SD (n=5/group). Significance levels: *P<0.05, **P<0.01, ***P<0.001, ****P<0.0001; ns=not significant.

### 3.3 Renal interstitial cell apoptosis was more likely to occur in the hyperglycemia group under water deprivation

Under water deprivation conditions, apoptosis was more pronounced in the high glucose group (Figure 2A,C). BAX expression was markedly elevated under high glucose conditions and further enhanced following water deprivation (Figure 2B,D). Conversely, BCL-2 expression was reduced in the high glucose environment (Figure 2B,E). The DM+WD group exhibited significantly increased cleaved Caspase-3 expression (Figure 2F,G). These findings indicate that under hyperglycemic conditions, water deprivation promotes renal cell apoptosis, potentially contributing to aggravated kidney injury in the high glucose state.

**Figure 2.**
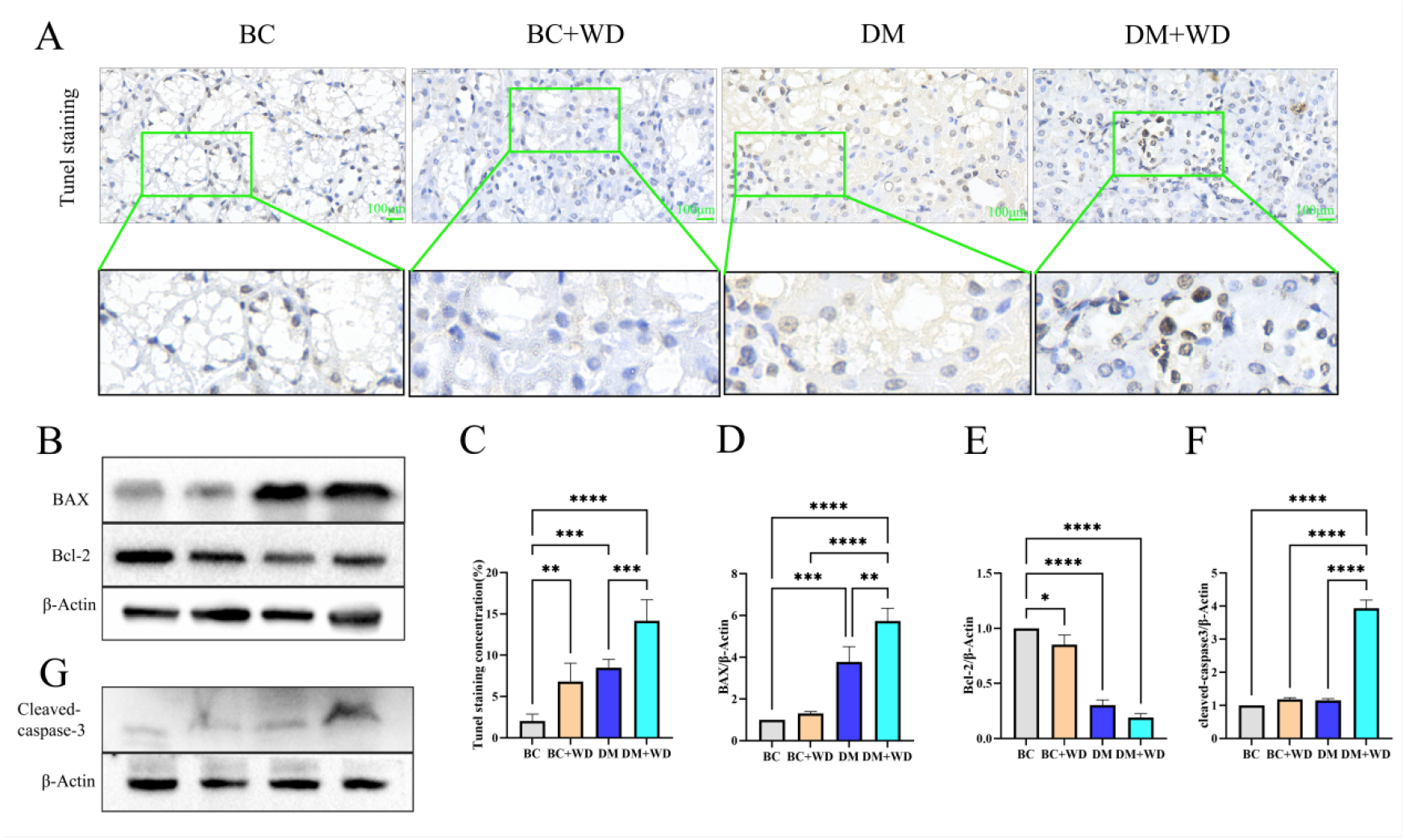
Enhanced renal apoptosis in diabetic mice under water deprivation. (A) TUNEL staining revealed significantly more apoptotic cells (green) in the renal interstitium of DM+WD mice compared to other groups (scale bar=100μm). (B) WB analysis demonstrated upregulated BAX and downregulated BCL-2 in DM+WD kidneys. (C) increased apoptotic index. (D) elevated BAX/β-actin ratio. (E) reduced BCL-2/β-actin ratio. (F) enhanced cleaved Caspase-3 activation in DM+WD group. (G) WB analysis of the expression of cleaved Caspase-3. with significance levels denoted by asterisks (*P<0.05, **P<0.01, ***P<0.001, ****P<0.0001) and ns indicating non-significance.

### 3.4 The expressions of gp91 and AQP2 enhanced after water deprivation challenge in a normal state, but this compensatory protective response was not obvious in DM

gp91(phox) (also known as NOX2) plays a crucial role in defending against infections while preventing excessive or chronic inflammation^[18-20]^. Our data revealed that water deprivation markedly elevated gp91 levels (indicated by green and yellow arrows), with more prominent expression observed in the BC+WD group (Figure 3A,C). This pattern was further confirmed by WB analysis (Figure 3B,D).

**Figure 3.**
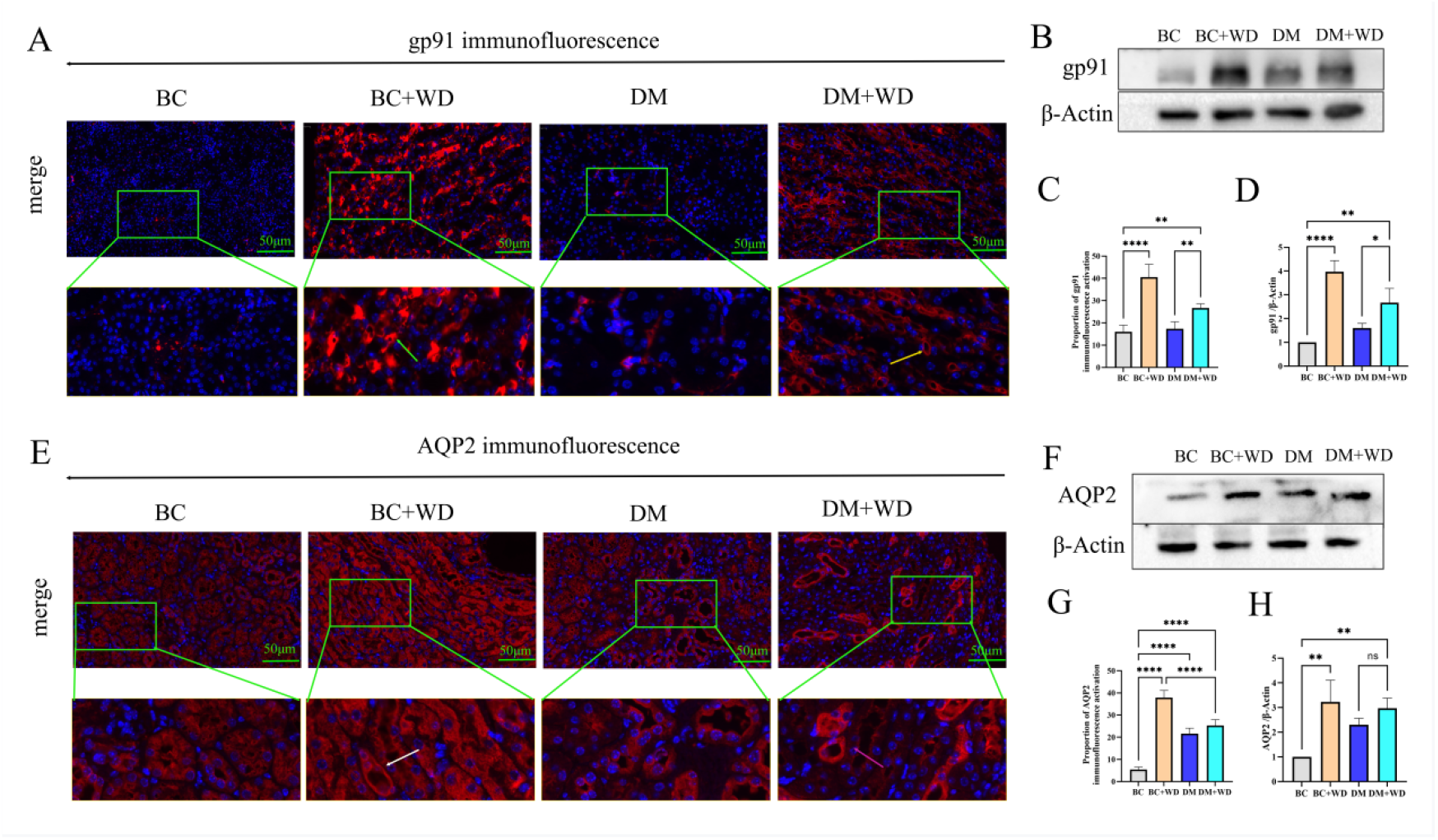
Blunted upregulation of renal gp91 and AQP2 in diabetic mice under water deprivation stress. (A) Immunofluorescence revealed stronger gp91 induction (green arrow) in renal tubules of BC+WD versus DM+WD mice (yellow arrows). (B) WB confirmed the diabetes-associated attenuation of gp91 upregulation. (C-D) Quantitative analyses demonstrated significantly lower gp91 induction in DM+WD kidneys compared to BC+WD controls. (E) Immunofluorescence revealed stronger AQP2 inductionin renal tubules of BC+WD (white arrow) versus DM+WD mice (pink arrows). (F) WB confirmed the diabetes-associated attenuation of AQP2 upregulation. (G,H) Quantitative analyses demonstrated significantly lower AQP2 induction in DM+WD kidneys compared to BC+WD. with significance levels denoted by asterisks (*P<0.05, **P<0.01, ***P<0.001, ****P<0.0001) and ns indicating non-significance.

AQP2, a pivotal protein in water balance regulation, demonstrated significantly increased expression following water deprivation in both BC+WD and DM+WD groups, as evidenced by immunofluorescence staining. However, the upregulation was more substantial in the BC+WD group (Figure 3E,G). These findings were also supported by WB experiments (Figure 3F,H).

Under physiological conditions, dehydration triggers a protective response in the kidney by upregulating gp91 and AQP2. However, in the diabetic state, although gp91 and AQP2 were still elevated, the additional elevation induced by water deprivation failed to provide sufficient renal protection to prevent injury.

### 3.5 FXR-TonEBP axis was enhanced after water deprivation challenge in a normal state, but this compensatory protective response blunted in DM

Immunofluorescence analysis revealed that water deprivation significantly increased FXR expression in the control water deprivation group, whereas this upregulation was less pronounced in the DM water deprivation group (Figure 4A,C). WB results confirmed the same trend in FXR expression before and after water deprivation (Figure 4B,D).

**Figure 4.**
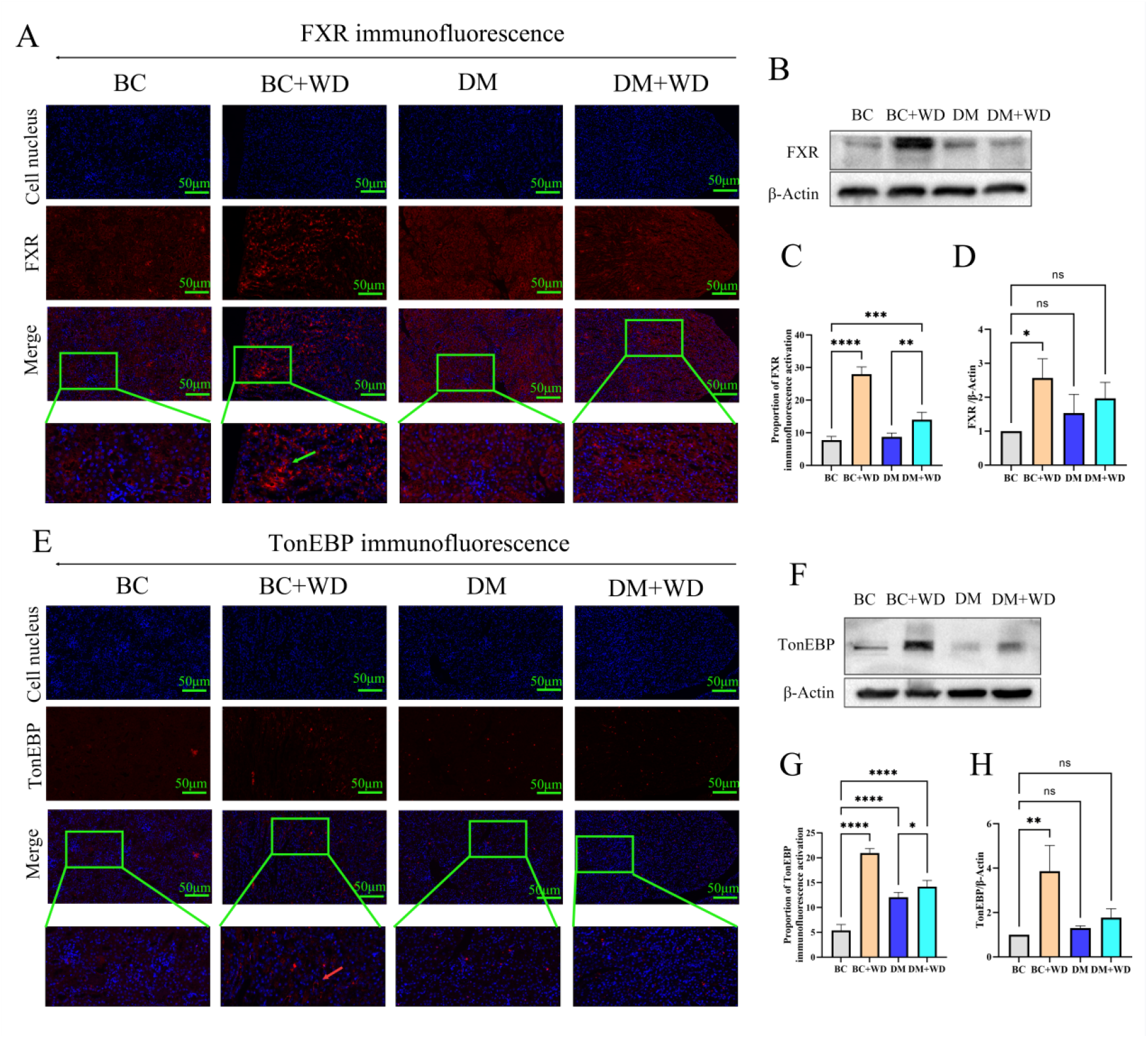
Impaired activation of the renal FXR-TonEBP axis in diabetic mice under water deprivation. (A) Immunofluorescence analysis revealed significantly enhanced nuclear FXR expression (green) in the renal tubules of WD, while DM mice showed attenuated induction (scale bar=50μm). (B) WB quantification confirmed the blunted FXR upregulation in DM+WD kidneys compared to controls. (C,D) Quantitative analyses demonstrated significantly lower FXR induction in DM+WD kidneys compared to BC+WD controls. (E) Immunofluorescence revealed stronger TonEBP inductionin renal tubules of BC+WD (red arrow) versus DM+WD mice.(F) WB quantification confirmed the blunted TonEBP upregulation in DM+WD kidneys compared to controls.(G,H) Quantitative analyses demonstrated significantly lower TonEBP induction in DM+WD kidneys.All quantitative data are shown as mean ± SD, *P<0.05, **P<0.01, ***P<0.001, ****P<0.0001, ns denotes statistically non-significant results.

Similarly, immunofluorescence demonstrated higher TonEBP expression in the control water deprivation group compared to the untreated control (Figure 4E,G), which was further validated by WB (Figure 4F,H). Both immunofluorescence and WB analyses indicated that TonEBP and FXR exhibited a weaker response to water deprivation in high-glucose conditions compared to non-DM mice.

The diminished FXR-TonEBP response under high glucose aligned with the reduced gp91/AQP2 expression. Impaired AQP2 upregulation in the high-glucose environment compromised renal medullary water reabsorption, leading to delayed kidney protection, increased urine output, and subsequent renal injury.

## 4. Discussion

Retrospective clinical analysis revealed that dehydration-associated kidney injury predominantly occurs in patients with preexisting conditions, with diabetic patients showing the worst renal prognosis. Animal experiments using intermittent water deprivation DM mice exhibited higher urine output and creatinine levels than non-diabetic controls, consistent with clinical observations. Renal pathology in DM mice after 16 weeks of intermittent dehydration showed significant damage to the medullary collecting ducts. Western blot and immunofluorescence analyses revealed attenuated expression of TonEBP and FXR in diabetic kidney tissue responding to dehydration compared to non-diabetic mice. This impairment further suppressed AQP2 expression, compromising water reabsorption and fluid balance, ultimately leading to renal interstitial cell apoptosis.

This investigation primarily examines how dehydration affects renal function under hyperglycemic conditions. The osmotic diuresis induced by elevated blood glucose levels results in significant water loss through urinary glucose excretion, leading to systemic dehydration that triggers thirst responses. While compensatory increases in water intake may temporarily relieve thirst and result in polyuria and polydipsia, inadequate fluid replacement during dehydration exacerbates the condition in diabetic individuals. This progressive dehydration ultimately causes structural and functional damage to the renal medulla^[21]^. Following rehydration, the impaired functional recovery of renal medulla leads to declined concentrating capacity, manifested as significantly increased urine output in diabetic mice after water restoration - a characteristic manifestation of DN development^[21, 22]^.Chronic severe dehydration in diabetic mice exacerbates renal injury, characterized by prominent interstitial fibrosis, impaired metabolic waste filtration, and consequent creatinine elevation^[23]^.

The pathophysiology of dehydration-induced renal injury involves three well-characterized mechanisms: (1) vasopressin-mediated hypertonic stress, (2) aldose reductase-fructokinase pathway activation, and (3) chronic hyperuricemia. These pathways collectively establish even mild dehydration as a modifiable risk factor for progression across chronic kidney disease subtypes^[24]^. Importantly, our study reveals a bidirectional relationship between dehydration and diabetes: while dehydration impairs pancreatic islet function (elevating blood glucose)^[25]^, the resulting hyperglycemia further exacerbates renal injury through increased glomerular filtration rate and impaired osmotic adaptation. This vicious cycle positions hyperglycemia not merely as a consequence, but as a critical amplifier of dehydration-induced kidney damage—a paradigm substantiated by our findings of blunted FXR-TonEBP activation in diabetic mice under dehydration stress.

TonEBP, predominantly localized in the renal medulla, serves as a critical transcriptional regulator of osmotic adaptation. Its nucleocytoplasmic shuttling responds dynamically to osmotic changes, coordinating the expression of genes encoding osmolyte transporters and synthetases that maintain intracellular ionic equilibrium under hypertonic stress ^[16, 26, 27]^. The NF-κB/FXR pathway exerts upstream control over TonEBP activation^[14]^, while downstream, TonEBP regulates water balance through aquaporin modulation - particularly AQP2-mediated water reabsorption^[11, 28]^. Through these dual mechanisms, TonEBP maintains medullary hypertonicity and urinary concentrating capacity, with its functional impairment leading to renal medullary atrophy^[29]^.

Our findings reveal a glucose-specific disruption of this protective system: diabetic kidneys show markedly attenuated activation of both FXR and TonEBP following dehydration. This impaired FXR-TonEBP axis directly compromises renal concentrating ability ^[13, 30]^ while concurrent dysregulation of NOX2/gp91 reflects broader cellular stress response alterations. The resulting downregulation of AQP2 and key metabolic effectors demonstrates a fundamental failure of osmotic adaptation in hyperglycemic conditions. Although ROS-mediated FXR suppression has been implicated in metabolic disorders ^[31]^, our study suggests that dehydration stress may be the background of glucose-induced FXR-tonebp pathway dysfunction, providing a new mechanistic basis for the progression of diabetic nephropathy^[32]^.

The impaired TonEBP-mediated response to dehydration under hyperglycemic conditions compromises renal hypertonic protection, promoting cellular apoptosis. Concurrently, suppressed FXR expression upregulates pro-apoptotic BAX while downregulating anti-apoptotic BCL-2, ultimately enhancing Cleaved caspase-3 activation^[33, 34]^, This apoptotic process further impairs renal water reabsorption capacity, representing a key pathogenic mechanism underlying diabetic polyuria. (**Figure 5**).

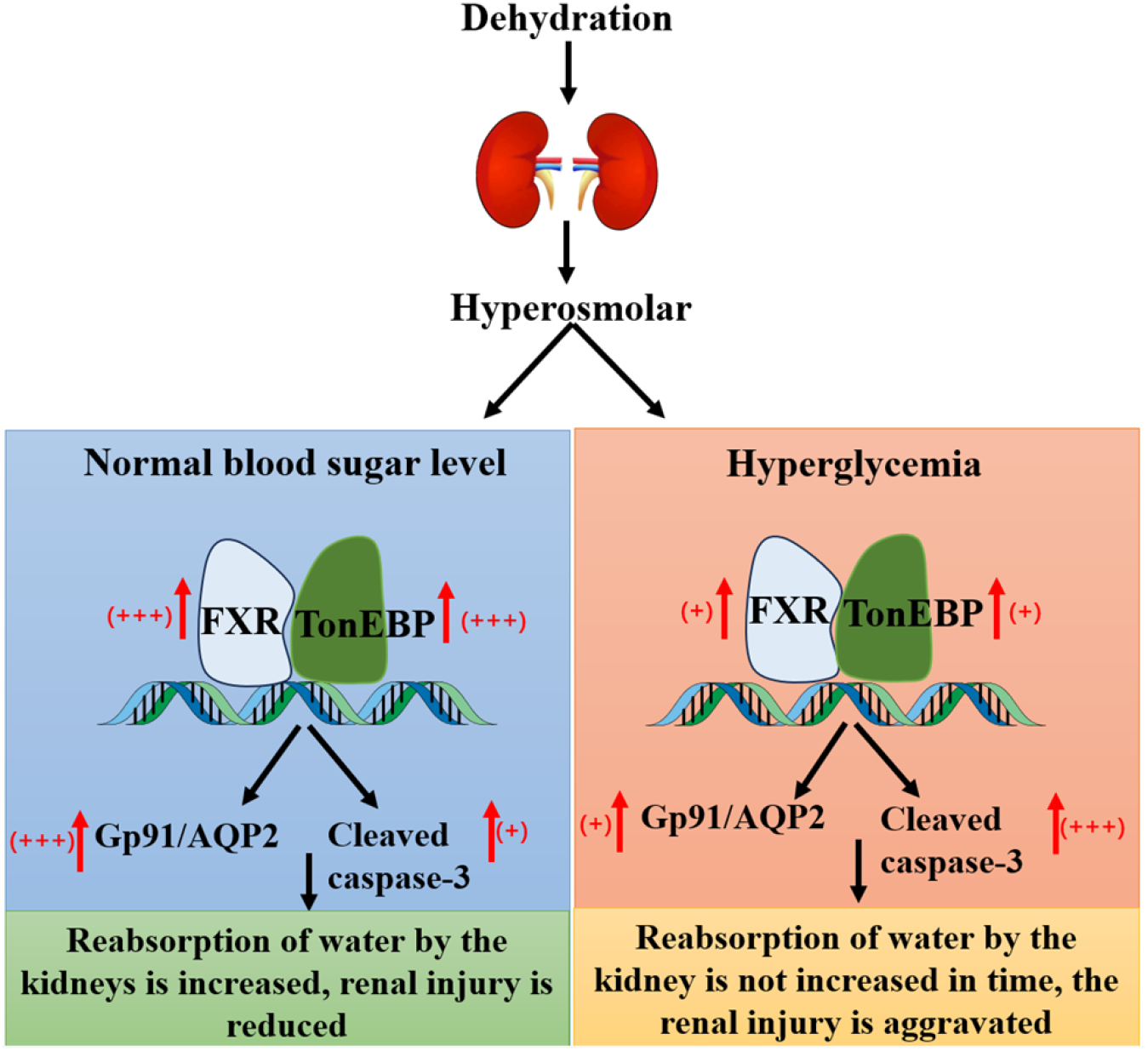

## Conclusion

In summary, this study demonstrates that water deprivation exacerbates renal injury in chronic conditions like hypertension and diabetes, with diabetic patients showing the poorest outcomes. The underlying mechanism involves glucose-impaired FXR-TonEBP signaling, which attenuates gp91/AQP2 expression and disrupts medullary osmotic regulation, ultimately inducing renal medullary cell necrosis. While these findings provide novel insights into diabetic nephropathy progression, certain limitations should be noted: the clinical data were retrospective in nature, and the animal model focused on type 1 diabetes, potentially limiting direct translation to type 2 diabetic nephropathy. Furthermore, the exact molecular link between hyperglycemia and FXR-TonEBP suppression warrants further investigation through targeted intervention studies. These limitations notwithstanding, our results establish a crucial role for osmotic stress dysregulation in diabetes-associated renal vulnerability.

## List of abbreviations

DM: diabetes mellitus
DN: diabetic nephropathy
FXR: farnesoid X receptor transcription factor
AQP2: aquaporin 2
gp91: glycoprotein 91
TonEBP: tonicity-responsive enhancer-binding protein
MCDs: medullary collecting duct cells
C57BL/6: C57 black 6
STZ: streptozotocin
SDS: sodium dodecyl sulfate
PVDF: polyvinylidene fluoride
HRP: horseradish peroxidase
PAS: Periodic Acid Schiff
PBST: phosphate buffered solution
Caspase-3: Cysteinyl aspartate specific protease-3

## Declarations

### Ethics approval

All animal experiments were approved by the Institutional Animal Care and Use Committee of Pukou Hospital of Traditional Chinese Medicine, Nanjing, China (Ethics Approval No. 20210026) and conducted in strict compliance with national guidelines for laboratory animal welfare. The study adheres to the ARRIVE guidelines for reporting animal research.

### Authors’ contributions

Tuo Wei,Liru Yin,and Jie Bo Huang conducted molecular genetic studies and sequence alignment, and drafted the manuscript. Fang Tian,Yan Li provided experimental design guidance. Qiong Wang and Enchao Zhou conceived the study, participated in its design and coordination, and assisted in manuscript preparation.

## Acknowledgements

We gratefully acknowledge the Central Laboratory of Jiangsu Provincial Hospital of Traditional Chinese Medicine and the Department of Basic Pharmacology for providing experimental facilities and technical support. A preliminary version of this work was previously shared as a preprint (https://www.researchsquare.com/article/rs-2308587/v1). The current manuscript includes expanded clinical data and new findings on FXR-TonEBP’s role in apoptosis and oxidative stress in water-deprived diabetic nephropathy mice, with all data and conclusions reflecting the present submission.

